# Identification of HCN1 as a 14-3-3 client

**DOI:** 10.1101/2021.08.19.457009

**Authors:** Colten Lankford, Jon Houtman, Sheila Baker

**Affiliations:** Department of Biochemistry and Molecular Biology, University of Iowa, Iowa City, IA 52242; Department of Microbiology and Immunology, University of Iowa, Iowa City, IA 52242

**Keywords:** 14-3-3 protein, ion channel, phosphorylation, hyperpolarization activated cyclic nucleotide gated channel (HCN1), protein turnover

## Abstract

Hyperpolarization activated cyclic nucleotide-gated channel 1 (HCN1) is expressed throughout the nervous system and is critical for regulating neuronal excitability, with mutations being associated with multiple forms of epilepsy. Adaptive modulation of HCN1 has been observed as has pathogenic dysregulation. While the mechanisms underlying this modulation remain incompletely understood, regulation of HCN1 has been shown to include phosphorylation. A candidate phosphorylation-dependent regulator of HCN1 channels is 14-3-3. We used bioinformatics to identify three potential 14-3-3 binding sites in HCN1. Isothermal titration calorimetry demonstrated that recombinant 14-3-3 binds all three phospho-peptides with low micromolar affinity. We confirmed that 14-3-3 could pull down HCN1 from multiple tissue sources and used HEK293 cells to detail the interaction. Two binding sites in the intrinsically disordered C-terminus of HCN1 were necessary and sufficient for a phosphorylation-dependent interaction with 14-3-3. The same region of HCN1 containing the 14-3-3 binding sites is required for phosphorylation-independent protein degradation. We propose a model in which phosphorylation of S810 and S867 (human S789 and S846) recruits 14-3-3 to inhibit a yet unidentified factor signaling for protein degradation, thus increasing the half-life of HCN1.

## Introduction

Hyperpolarization activated currents (*I*_h_) carried by HCN1 channels play essential roles throughout the nervous system in tempering excitability, setting pacemaking activity, and coordinating signal integration [1-3]. Mutations that prevent ion conduction or alter gating are associated with high seizure susceptibility which can result in an extremely limited lifespan. Mutations affecting regions outside of the pore forming and voltage sensing domains may result in more mild generalized epilepsy or intellectual disability [4-6]. Given its role controlling excitability, it is unsurprising to find that HCN1 is implicated in many neurological functions. HCN1 plays a complex role in learning and memory with global knockout of HCN1 resulting in motor learning deficits while knockout of HCN1 selectively from the forebrain enhanced spatial reference memory [7, 8]. HCN1 knockdown in the CA1 region was shown to be anxiolytic [9]. In the forebrain HCN1 is inhibited by ketamine, contributing to the unique profile of this anesthetic and psychotropic drug [10, 11]. HCN1 is required in the olfactory bulb for activity-dependent neurogenesis [12]. And in the retina, HCN1 is required to shape the output of rod photoreceptors to prevent saturation of inner retinal circuitry [13-15]. HCN1 is an attractive pharmaceutical target [16]. The diversity of processes in which HCN1 participates makes it essential to uncover mechanisms of differential regulation. Further, the similarity between HCN1 and other HCN channels, including HCN4 the primary driver of cardiac rhythmicity [17, 18], highlights the need to identify possible drug targets specific for HCN1.

HCN1 is not a static channel but can be dynamically regulated. In hippocampal CA1 neurons, dendritic HCN1 participates in non-synaptic plasticity where HCN1 activity is downregulated to enhance spike probability and upregulated to reduce excitability [7, 19]. In addition, HCN1 channels are modulated in a homeostatic manner to prevent excessive signaling by strengthened synapses [20]. The neuromodulators neuropeptide Y and corticotrophin-releasing factor were shown to differentially modulate HCN1 activity in pyramidal neurons of the basolateral amygdala, downregulating and upregulating HCN1 respectively [21]. Dysregulation of HCN1 has also been observed and is associated with pathological states. Both acute and chronic HCN1 downregulation was observed in the hippocampus of a rat model of status epilepticus, leading to the speculation that dysregulation of HCN1 contributes to this condition [22]. Changes in HCN1 activity are also implicated in neuropathic pain. In layer 5 pyramidal neurons of the anterior cingulate cortex HCN1 was downregulated under chronic constriction injury [23]. Conversely, HCN1 has been shown to be upregulated in the periphery in models of neuropathic pain and has been shown to be a therapeutic target [24-28].

The activity, trafficking, and stability of HCN1 channels can be regulated by the accessory subunit, TRIP8b. TRIP8b binds to the cytoplasmic C-terminal tail in a bimodal manner, making contact with the membrane proximal Cyclic-Nucleotide Binding Domain (CNBD) and the terminal tripeptide ser-asn-leu, thus bridging a 300 amino acid long intrinsically disordered region [29]. Differential regulation of HCN1 by TRIP8b can be accomplished through the expression of different TRIP8b splice isoforms [30-32]. Additionally, TRIP8b can be phosphorylated by CaMKIIa or PKA at a site involved in binding to the CNBD [33]. Two additional HCN1 binding proteins are the actin bundling protein filamin A, and the ubiquitin ligase Nedd4-2, both of which can limit HCN1 surface expression [34, 35]. However, little is known about how the interaction between these proteins and HCN1 may be regulated. HCN1 itself is phosphorylated at multiple sites. Tyrosine phosphorylation or p38MAPK signaling can alter channel activity while PKC-dependent phosphorylation can alter surface expression [23, 36, 37].

Despite phosphorylation being implicated in HCN1 modulation, the effectors reading this post-translational modification are not known. 14-3-3 is a strong candidate for such an effector. 14-3-3 proteins are a conserved family of phosphoserine/phosphothreonine binding proteins ubiquitously expressed in eukaryotes and particularly enriched in the brain. There are over a hundred proteins that have been thoroughly validated as 14-3-3 client proteins. These include proteins participating in cellular functions as diverse as fatty acid metabolism and apoptosis [38, 39]. Bioinformatics predict the 14-3-3 ‘client-ome’ actually consists of several thousand proteins and is enriched in 2R-ohnologues [40]. HCN1 is an example of a 2R-ohnologue, which are small (two or four member) protein families arising from whole genome duplications. Among the 14-3-3 client proteins are multiple ion channels for which 14-3-3 may modulate channel assembly, trafficking, or gating [41-44]. These observations prompted us to investigate the possibility that 14-3-3 could be a phosphorylation-dependent regulator of HCN1.

Here we demonstrate that HCN1 is a novel 14-3-3 client protein and this interaction requires phosphorylation of two serines in the intrinsically disordered C-terminal tail of HCN1. Using cultured cells, we observed that HCN1 mutants lacking these phosphorylatable serine residues behaved identically to wild-type channels, but perturbations of the surrounding residues resulted in stabilization of HCN1 channels. The mechanism controlling degradation of HCN1 is unknown. Our data supports a model in which phosphorylation of serines 867 and 810 signals a switch in binding of a yet unidentified factor promoting degradation to binding of 14-3-3 which may act to stabilize HCN1 expression.

## Results

### Potential 14-3-3 binding sites in the C-terminus influence HCN1 expression in HEK293 cells

We analyzed the sequence of HCN1 using 14-3-3Pred, a prediction tool for the identification of phospho-peptides likely to serve as 14-3-3 ligands [45]. Several potential 14-3-3 binding sites (underlined, Fig. 1) were predicted. We limited our focus to unstructured regions as 14-3-3 would likely have limited accessibility to binding motifs within the structured regions such as the HCN domain and cyclic nucleotide binding domain. We elected to evaluate the four highest scoring peptides that are conserved in vertebrates (boxed, Fig. 1). All four sites are located in the long, intrinsically disordered portion of the HCN1 C-terminal tail which contains 37 serines and 30 threonines, 49% of all serines and threonines in HCN1. One of those peptides overlaps the terminal ser-asn-leu tripeptide required for binding to the accessory subunit TRIP8b thus would be inaccessible to 14-3-3 *in vivo* [29]. We focused on testing the other three peptides (A) ^647^SRMRTQ[**pS**]PPVY, (B) ^804^VRPLSA[**pS**]QPSL, and (C) ^861^TLFRQM[**pS**]SGAI using isothermal titration calorimetry (ITC). Previous studies of 14-3-3 using ITC demonstrate that most singly phosphorylated peptide ligands bind with a K_*D*_ in the low micromolar range of 0.1 - 60 µM [46-54]. All three HCN1 phosphopeptides were bound by purified 14-3-3ζ with similar affinity; 16 µM (peptide A), 25 µM (peptide B), and 15 µM (peptide C) (Fig. 2 and Table 1). This confirms that these three predicted serine-phosphorylated peptides from HCN1 can be bound by 14-3-3.

**Table 1:**
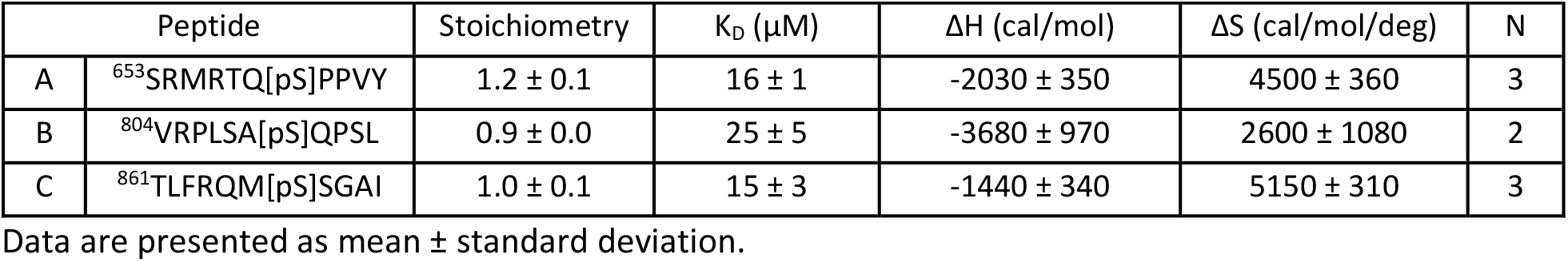
Expanded ITC data related to Figure 2

**Figure 1:**
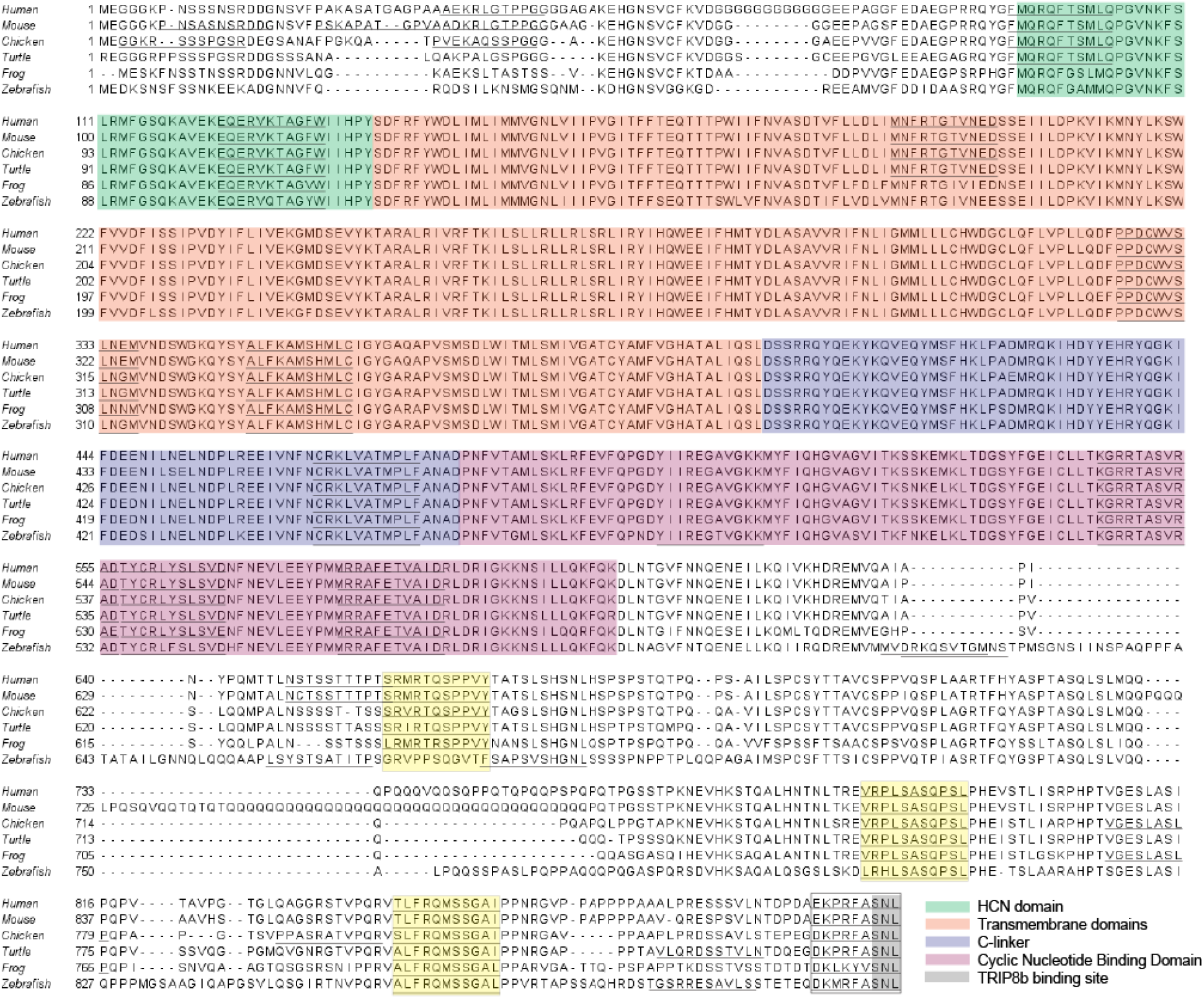
The C-terminus of HCN1 contains multiple predicted 14-3-3 binding sites. Sequence alignment of HCN1 from five vertebrate species. 14-3-3Pred predicted 14-3-3 binding sites are underlined. The structured domains of HCN1 highlighted as follows: green = HCN domain, orange = transmembrane domains, blue = C-linker, and purple = Cyclic Nucleotide Binding Domain). Conserved 14-3-3 binding motifs in exposed cytoplasmic domains are boxed and the three sites of focus are shaded in yellow.

**Figure 2:**
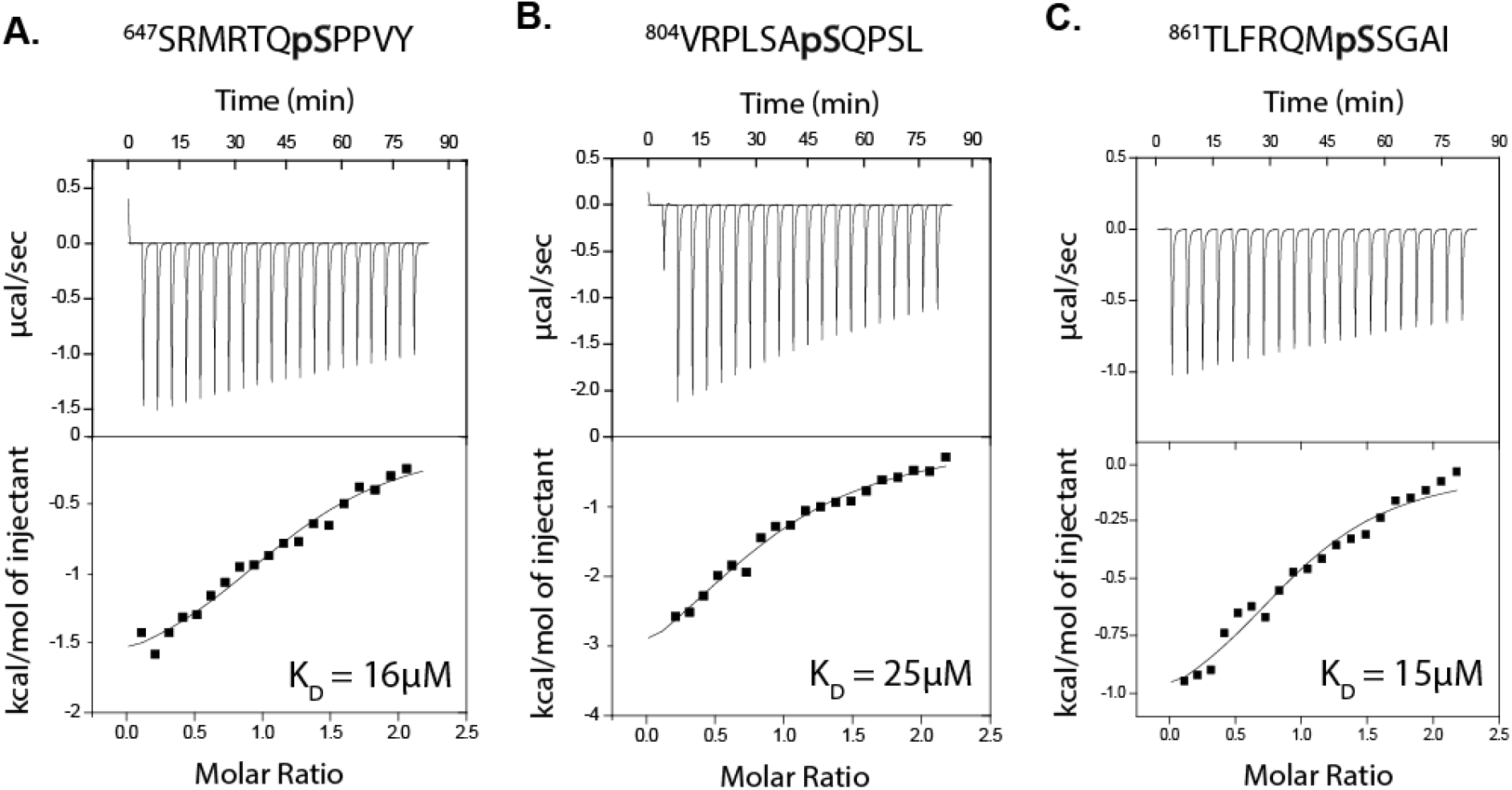
14-3-3 binds to phosphopeptides corresponding to HCN1 sequences. Representative isothermal titration (ITC) binding curve with the average dissociation constant (K_D_) noted for phospho-peptides corresponding to 14-3-3 binding site **A, B**, and **C**. Additional details are provided in Table 1.

To examine whether 14-3-3 could potentially regulate HCN1 activity, we generated stably integrated HEK293 cell lines expressing either mRuby3-HCN1 or mRuby3-HCN1_Δ3,14-3-3_. In mRuby3-HCN1_Δ3,14-3-3_ the amino acids corresponding to 14-3-3 binding peptides B and C (residues 804-814 and 861-871) were deleted while targeted alanine substitution was performed at the 14-3-3 binding site corresponding to peptide A to prevent phosphorylation while preserving a previously described di-arginine ER retention motif (^647^S<u>R</u>M<u>R</u>**T**Q**S**PPVY to ^647^S<u>R</u>M<u>R</u>**A**Q**A**PPVY) [55]. Whole-cell current recordings from these two cell lines revealed that the activation voltage of the mutant HCN1 was similar to the wild-type; mRuby3-HCN1 V_1/2_: -118.6 ± 0.1 mV and mRuby3-HCN1_Δ3, 14-3-3_ V_1/2_: -116.1 ± 0.1 mV (Fig. 3A). However, there was increased current density in cells expressing the mutant HCN1; maximal current density of mRuby3-HCN1: -34.64 ± 6.37 pA/pF vs mRuby3-HCN1_Δ3, 14-3-3_: -69.10 ± 10.34 pA/pF; p = 0.0047 (Fig. 3B, C). Complimentary surface biotinylation experiments revealed that mRuby3-HCN1_Δ3,14-3-3_ had higher total and surface expression levels compared to mRuby3-HCN1; total: 2.10 ± 0.30-fold increase compared to mRuby3-HCN1; p < 0.0001 and surface: 2.50 ± 0.10-fold increase compared to mRuby3-HCN1; p = 0.0071 (Fig. 3D, E). From these experiments we conclude that 14-3-3 does not regulate activation of the channel but is implicated in the turnover or trafficking of HCN1.

**Figure 3:**
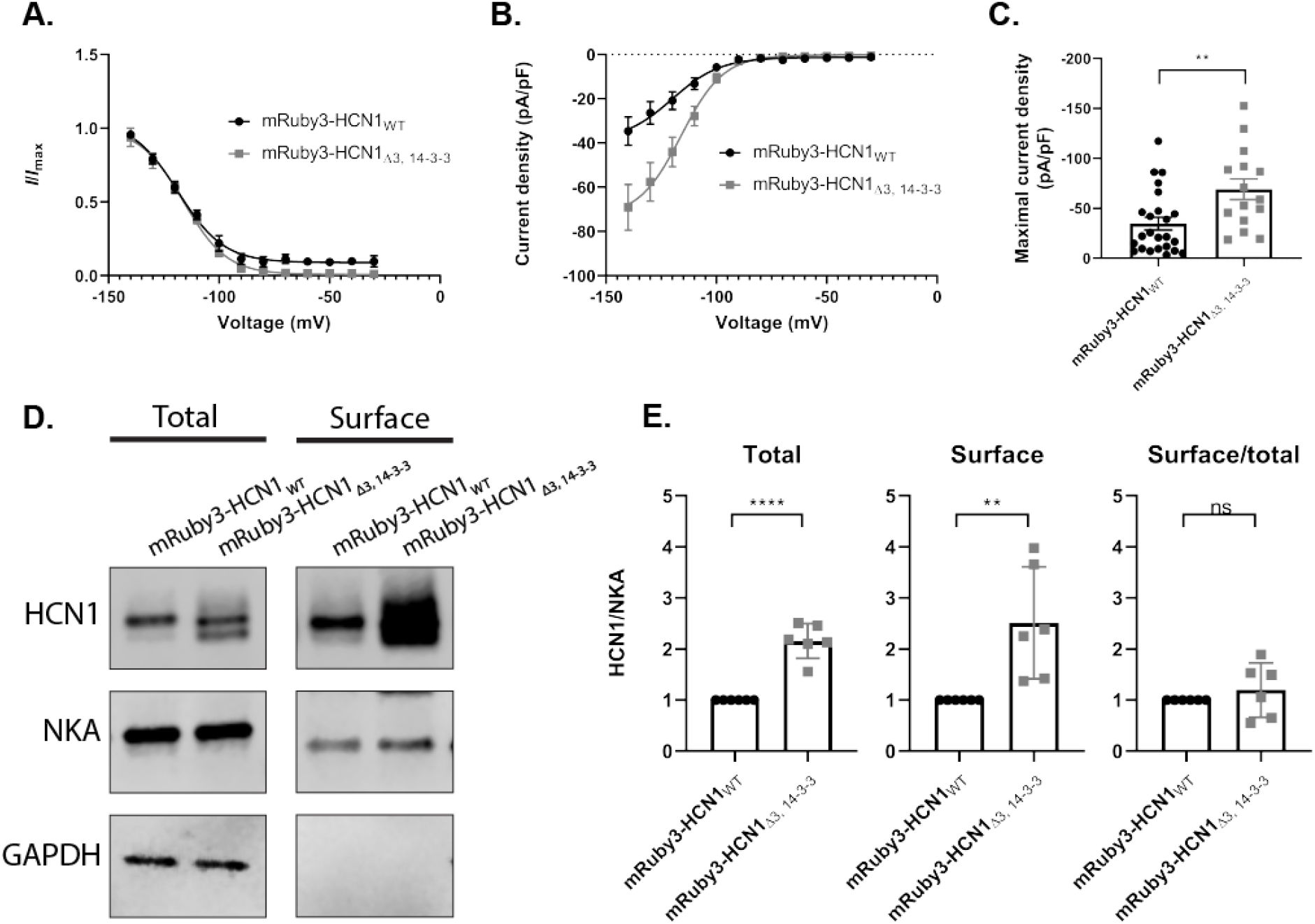
Mutating 14-3-3 binding sites enhances HCN1 protein level in stable HEK293 cell lines. Activation voltage **(A)**, current density **(B)**, and maximal current density of *I*_h_ **(C)**, recorded from HEK293 cell lines expressing mRuby3-HCN1 (black) or mRuby3-HCN1_Δ3, 14-3-3_ (grey). **D)** Representative western blot showing total HCN1 expression and surface HCN1 expression isolated by surface biotinylation. Western blotting against the endogenous NKA and GAPDH as membrane and cytosolic protein controls. **E)** Relative amount of total and surface mRuby3-HCN1_Δ3, 14-3-3_ compared to mRuby3-HCN1 normalized to NKA. Error bars represent standard error of mean for electrophysiology data (A, B, C) and standard deviation for western blot data (E). T-test p < 0.05 (*); p < 0.01 (**); p<0.001 (***); p < 0.0001 (****).

### 14-3-3 interacts with HCN1

To determine if HCN1 and 14-3-3 interact, we used a pan-14-3-3 antibody to immunoprecipitate 14-3-3 from retina lysates. HCN1 was co-immunoprecipitated from wild-type retina but not HCN1 knockout retina which was used to ensure specificity of the HCN1 signal. An anti-GFP antibody was used as a negative control to ensure HCN1 was not binding non-specifically to the beads or mouse IgG (Fig. 4A). Next, we performed pull-down assays using recombinant _6His_14-3-3ζ. _6His_14-3-3ζ immobilized on Ni-NTA beads, but not the unconjugated control beads, pulled down HCN1 from whole brain lysate, retinal lysate, and lysate from HCN1 expressing HEK293 cells (Fig 4B). We conclude that HCN1 can interact with 14-3-3 although it is not possible from these assays to determine if this interaction is direct.

**Figure 4:**
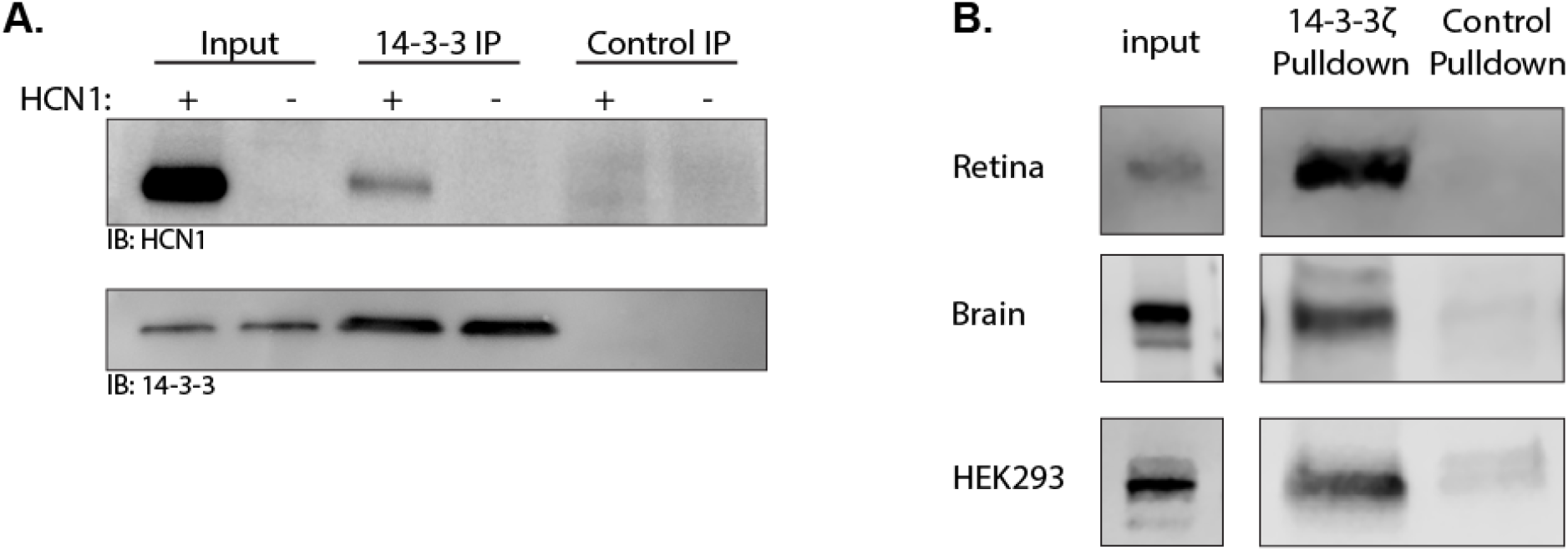
14-3-3 interacts with HCN1. **A)** Western blotting for HCN1 and 14-3-3 following 14-3-3 IP or control GFP IP from wild-type and HCN1 KO retina. **B)** Western blotting for HCN1 following pulldown with recombinant 14-3-3ζ or resin only control from mouse retina, whole brain, and HCN1 expressing HEK293 cells.

### 14-3-3 binds two sites at the distal C-terminus of HCN1

To determine if all three 14-3-3 binding sites are used in living cells, and to assay for any additional sites 14-3-3Pred may not have scored highly, we used transiently transfected HEK293 cell lysates as a source of HCN1 protein fragments in pull down assays with immobilized _6His_14-3-3ζ. We first tested if the C-terminus of HCN1 was sufficient for 14-3-3 binding. Amino acids 598-910 of HCN1 (CT_598-910_) was fused to a transmembrane domain (from the PDGF-receptor) with an extracellular SNAP-tag as shown in Fig 5A. CT_598-910_, but not the control base reporter, was pulled down by 14-3-3 (Fig. 5C, D). Serial truncations of the HCN1 fragment demonstrated that loss of residues 803-910 (containing peptides B and C) severely limited the interaction (0.20 ± 0.10-fold decrease compared to CT_598-910_; p < 0.0001). Further deletion that left peptide A intact (CT_598-698_) resulted in complete loss of interaction (0.06 ± 0.04-fold decrease compared to CT_598-910_; p < 0.0001) (Fig 5C, D). The complimentary deletion of the peptide A-containing region confirmed it was not necessary for 14-3-3 binding (Fig 5G). It may be that site A is not used *in vivo* at all, or simply that it was not phosphorylated in this model system.

**Figure 5:**
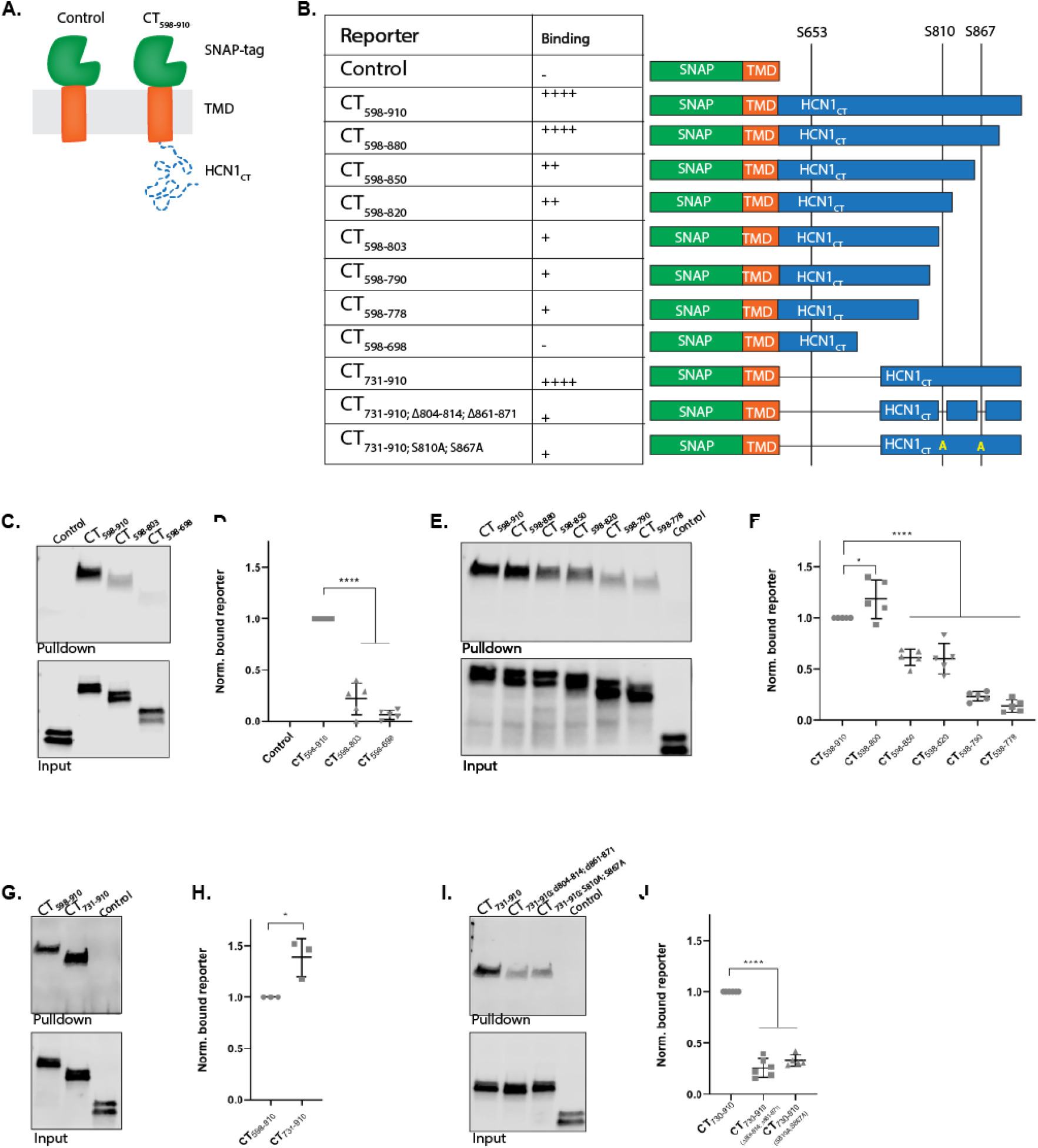
14-3-3 binds to two sites in the distal C-terminus of HCN1. **A)** Schematic of the reporter system used in this series of experiments. The C-terminus of HCN1 (residues 598-910) was fused to the transmembrane reporter. Reporters were expressed in HEK293 cells and pulled down with recombinant 14-3-3ζ then analyzed by western blotting for the SNAP-tag. **B)** Summary of reporters used. Binding is given as efficiency compared to full-length HCN1 C-terminus with ++++ = 100%, +++ = 75%, ++ = 50%, + = 25%, and - = 0% binding efficiency. **C, E, G, I)** Representative blots and comparisons of normalized results **(D, F, H, J)** for pulldowns of the reporter-HCN_CT_ constructs as indicated. For each construct, the p-value or adjusted p-value for comparison to CT_598-910_ or CT_731-910_ is shown p < 0.05 (*); p < 0.01 (**); p<0.001 (***); p < 0.0001 (****).

Serial deletions of 30 amino acid blocks from the most distal portion of the C-terminal tail of HCN1 generated three findings (Fig. 5E, F). First, deleting the block containing the TRIP8b-tripeptide binding site did not impede the interaction as slightly more of this truncated reporter was pulled down (1.18 ± 0.18-fold increase compared to CT_598-910_; p = 0.0359), supporting our earlier decision to disregard the prediction of a fourth putative 14-3-3 binding site. Second, HCN1 fragments lacking the region containing peptide C (amino acids 850-880) resulted in loss of half the binding efficiency (0.61 ± 0.07-fold decrease compared to CT_598-910_; p < 0.0001). Third, another drop in binding efficiency was observed when the serial truncation also resulted in loss of the region containing peptide B (residues 790-820; 0.23 ± 0.04-fold decrease compared to CT_598-910_; p < 0.0001).

To determine whether the distalmost C-terminus of HCN1 was sufficient to mediate the interaction with 14-3-3, the proximal region of the C-terminus including peptide A was removed leaving only a 180 amino acid long fragment of the distal C-terminus that retains the disordered polyQ stretch as a spatial linker (residues 731-910). This construct was pulled down slightly more efficiently than the construct containing the full-length C-terminus (Fig. 5G, H; 1.39 ± 0.19-fold increase compared to CT_598-910_; p = 0.0227). Finally, site-specific mutations were used to delete just the 10 amino acid blocks corresponding to peptides B and C from the distal C-terminus (CT_731-910; Δ804-814; Δ861-871_) or to mutate the phosphorylation sites of those peptides with serine to alanine replacements (CT_731-910; S810A; S867A_). Both sets of mutations reduced 14-3-3 binding significantly (Fig. 5I, J; 0.26 ± 0.09-fold decrease compared to CT_731-910_ for CT_731-910; Δ804-814; Δ861-871_ and 0.33 ± 0.05-fold CT_731-910_ for CT_731-910S; S810A; S867A_; p < 0.0001 for both). This data indicates that 14-3-3 binds to two sites in the disordered tail of HCN1 centered around S810 and S897.

### Kinase activity is required for interaction between 14-3-3 and HCN1

We next tested which kinase could be phosphorylating S810 and/or S897 to allow 14-3-3 recruitment. HEK293 cells transfected with the reporter CT_731-910_ were treated with a panel of kinase inhibitors (detailed in Supplementary Table 1) for three hours, then tested for interaction with 14-3-3 using the pull-down assay. Treatment with the broad-spectrum kinase inhibitor, staurosporine reduced the efficiency of CT_731-910_ pulldown similar to the S810A; S867A mutation (Fig. 6; 0.26 ± 0.10-fold decrease compared to untreated control; p < 0.0001). Treatment with more specific kinase inhibitors had lesser effects and generally any single inhibitor slightly reduced the amount of CT_731-910_ pulled down. Only treatment with the PKC inhibitor (bisindolylmaleimide I) and the p38MAPK inhibitor (PD 169316) had a statistically significant effect on the amount of reporter pulled down by _6His_14-3-3ζ (PKC inhibition: 0.61 ± 0.11-fold decrease compared to untreated control; p = 0.0468 and p38MAPK inhibition: 0.61 ± 0.19-fold decrease compared to untreated control; p = 0.0468). The need for kinase activity is consistent with the canonical mechanism of 14-3-3 binding.

**Figure 6:**
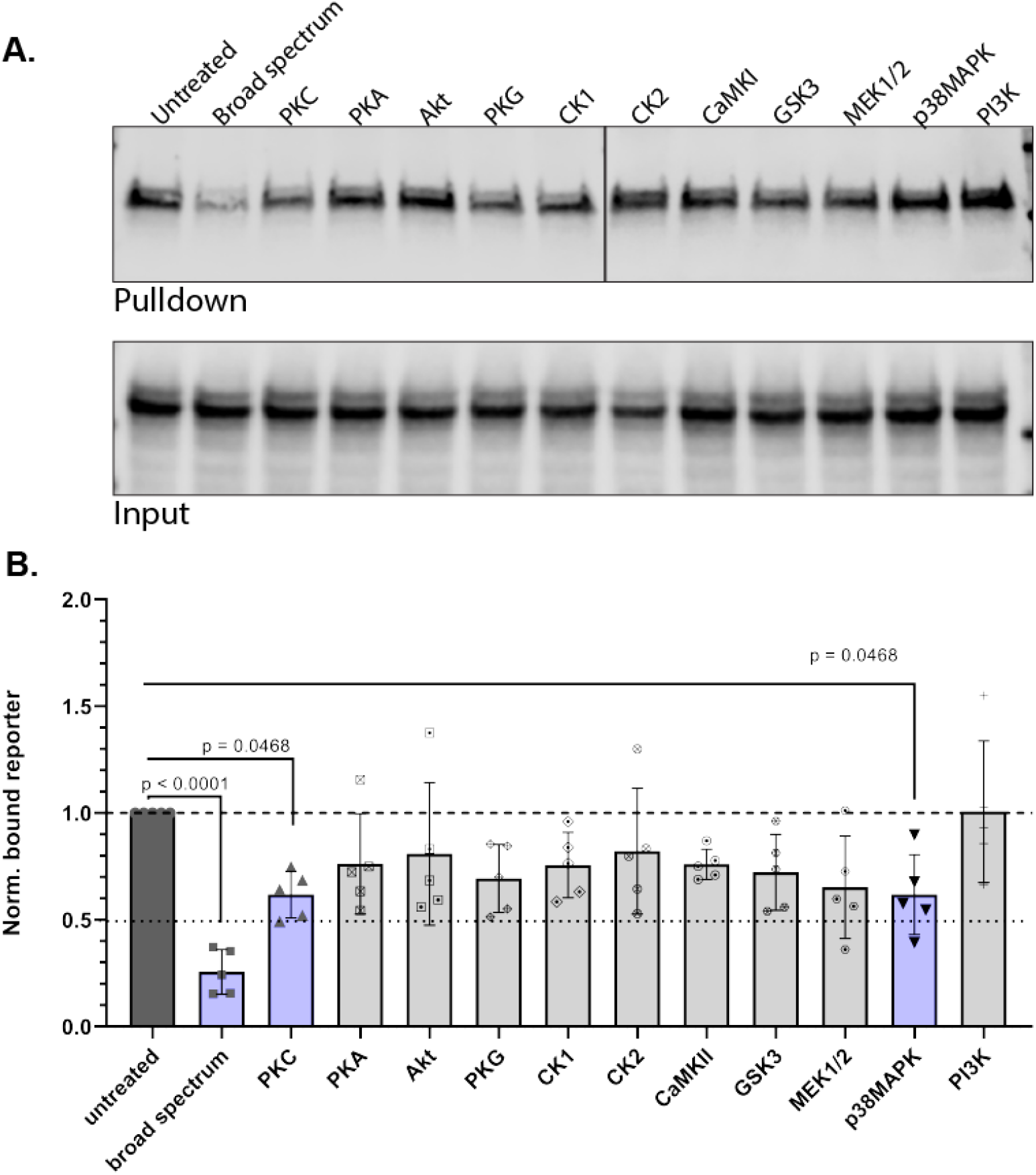
14-3-3 binding requires HCN1 phosphorylation. HEK293 cells expressing reporter-HCN1 CT_731-910_ were treated with kinase inhibitors (detailed in Table S1) **A)** representative pulldown **B)** comparison of all experimental replicates, dark grey bar is the untreated control, purple bars highlight the treatments that generated a statically significant reduction in pulldown efficiency with adjusted p-value shown.

### 14-3-3 binds to a region that negatively regulates HCN1 expression

Having determined that only two of the initially tested HCN1 phosphopeptides are involved in 14-3-3 binding in HEK293 cells we tested the effect of more specific mutations on the expression level of HCN1. Stable cell lines were generated for expression of SNAP-tagged full-length HCN1 and two mutant versions of HCN1 – one with both 14-3-3 binding sites entirely deleted (SNAP-HCN1_Δ804-814; Δ861-871_) and the other with just the central serine residues replaced with alanine (SNAP-HCN1_S810A; S867A_). HCN1 expression was measured in surface biotinylation assays. We expected both mutant versions of HCN1 would be expressed at ∼2 fold higher levels as we saw previously for HCN1_Δ3, 14-3-3_. However, SNAP-HCN1_Δ804-814; Δ861-871_ expressed at roughly six-times the level of SNAP-HCN1 both in total (6.17 ± 1.20-fold increase compared to SNAP-HCN1; p = 0.0003) and at the surface (5.75 ± 0.08-fold increase compared to SNAP-HCN1; p < 0.0001). To our surprise, SNAP-HCN1_S810A; S867A_ did not exhibit a change in total expression and the change in surface expression was small (1.36 ± 0.09-fold increase compared to SNAP-HCN1; p = 0.0018) and unlikely to be biologically significant (Fig. 7A, B).

**Figure 7:**
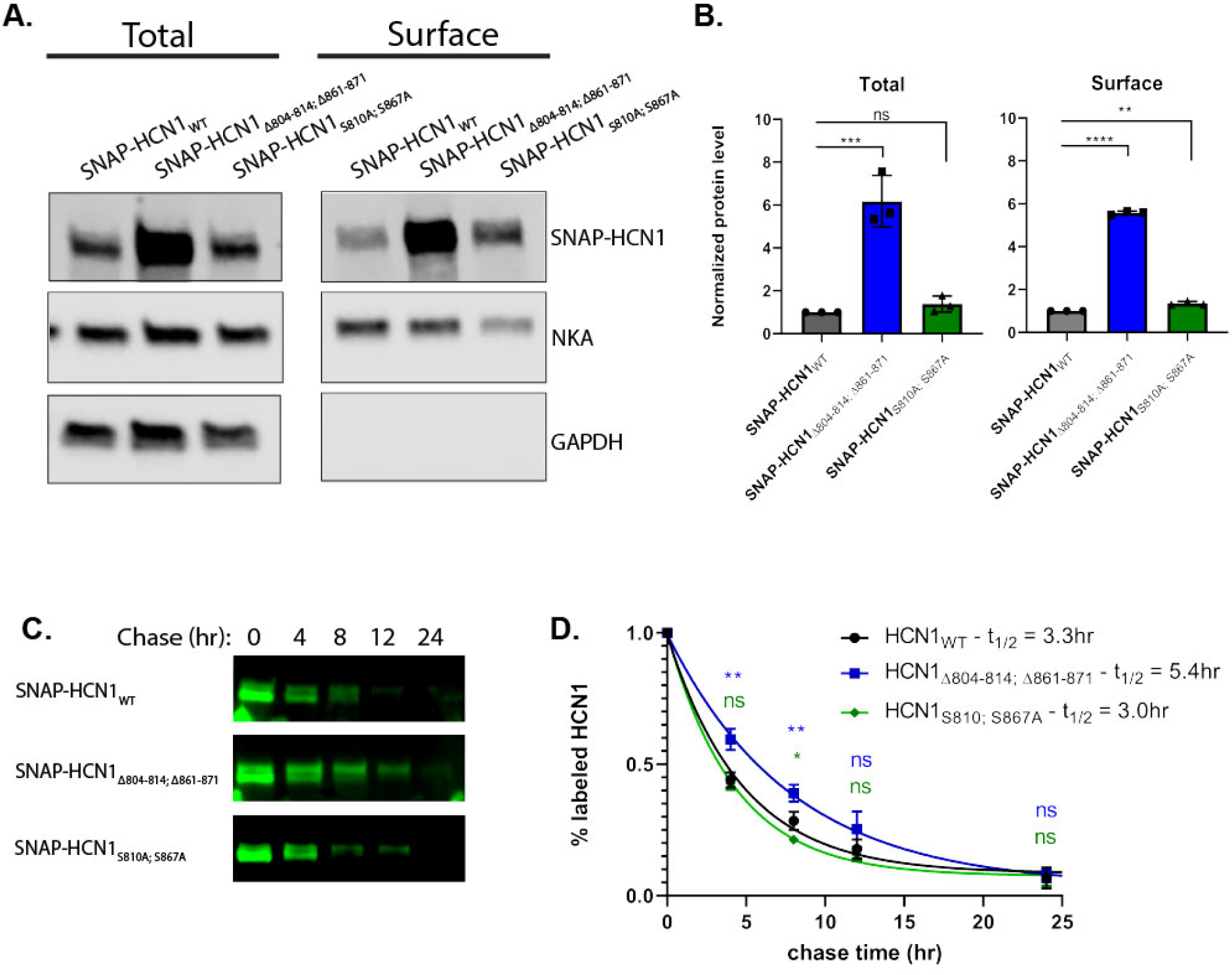
14-3-3 binding sites regulate HCN1 turnover in a phosphorylation independent manner. **A)** Representative western blot showing total SNAP-HCN1 expression and surface SNAP-HCN1 expression isolated by surface biotinylation with western blotting against the endogenous NKA and GAPDH as membrane and cytosolic protein controls. B) Relative total and surface level of SNAP-HCN1_Δ804-814; Δ861-871_ (blue) and SNAP-HCN1_S810A; S867A_ (green) compared to SNAP-HCN1 (black) normalized to NKA. **C)** Representative fluorescence imaging of labeled SNAP-HCN1 harvested after 0, 4, 8, 12, and 24 hours of chase. **D)** Relative amount of labeled SNAP-HCN1 (black), SNAP-HCN1_Δ804-814; Δ861-871_ (blue), and SNAP-HCN1_S810A; S867A_ (green), normalized to amount at 0 hour chase plotted against chase time to give a decay curve. The adjusted p-value for comparison to SNAP-HCN1 is shown p < 0.05 (*); p < 0.01 (**); p<0.001 (***); p < 0.0001 (****).

To investigate the dynamics of SNAP-HCN1 expression we used pulse-chase assays exploiting the ability of the SNAP-tag to covalently bind substrate. Cells were incubated with a fluorescent SNAP-substrate for 45 min which was then washed out and replaced with a non-fluorescent SNAP-substrate so that any SNAP-HCN1 synthesized after the pulse could not be fluorescently labeled. Cells were harvested at several time points for the next 24 hours. Following SDS-PAGE and transfer onto PVDF membrane, the amount of labeled SNAP-HCN1 was measured by direct fluorescence detection and the amount of total SNAP-HCN1 was measured by western blotting against the SNAP-tag (Fig. 7C). The half-life was calculated by fitting the normalized data to decay curves (Fig. 7D). The half-life of SNAP-HCN1 was 3.3 hours. The half-life of SNAP-HCN1_S810A; S867A_ was slightly reduced to 3.0 hours and the relative amount of labeled SNAP-HCN1_S810A; S867A_ was decreased slightly compared to SNAP-HCN1 at 8 hours (0.21 ± 0.02 vs 0.28 ± 0.03; p = 0.0278). However, the half-life of SNAP-HCN1_Δ804-814; Δ861-871_ was significantly increased to 5.4 hours with the relative amount of labeled SNAP-HCN1_Δ804-814; Δ861-871_ being elevated compared to SNAP-HCN1 at 4 hours (0.59 ± 0.04 vs 0.44 ± 0.03; p = 0.0021) and 8 hours (0.39 ± 0.03 vs 0.28 ± 0.03; p = 0.0077). A longer half-life indicates increased stability of the protein and would lead to the increased static expression levels we observed.

In conclusion, we document that HCN1 is a client of 14-3-3. In HEK293 cells, kinase activity and serine at two different 14-3-3 binding sites within the distal C-terminal tail of HCN1 is required for the interaction. Our data also indicate that the region of HCN1 involved in 14-3-3 interaction overlaps with a phosphorylation independent regulator of HCN1 turnover (Fig. 8).

**Figure 8:**
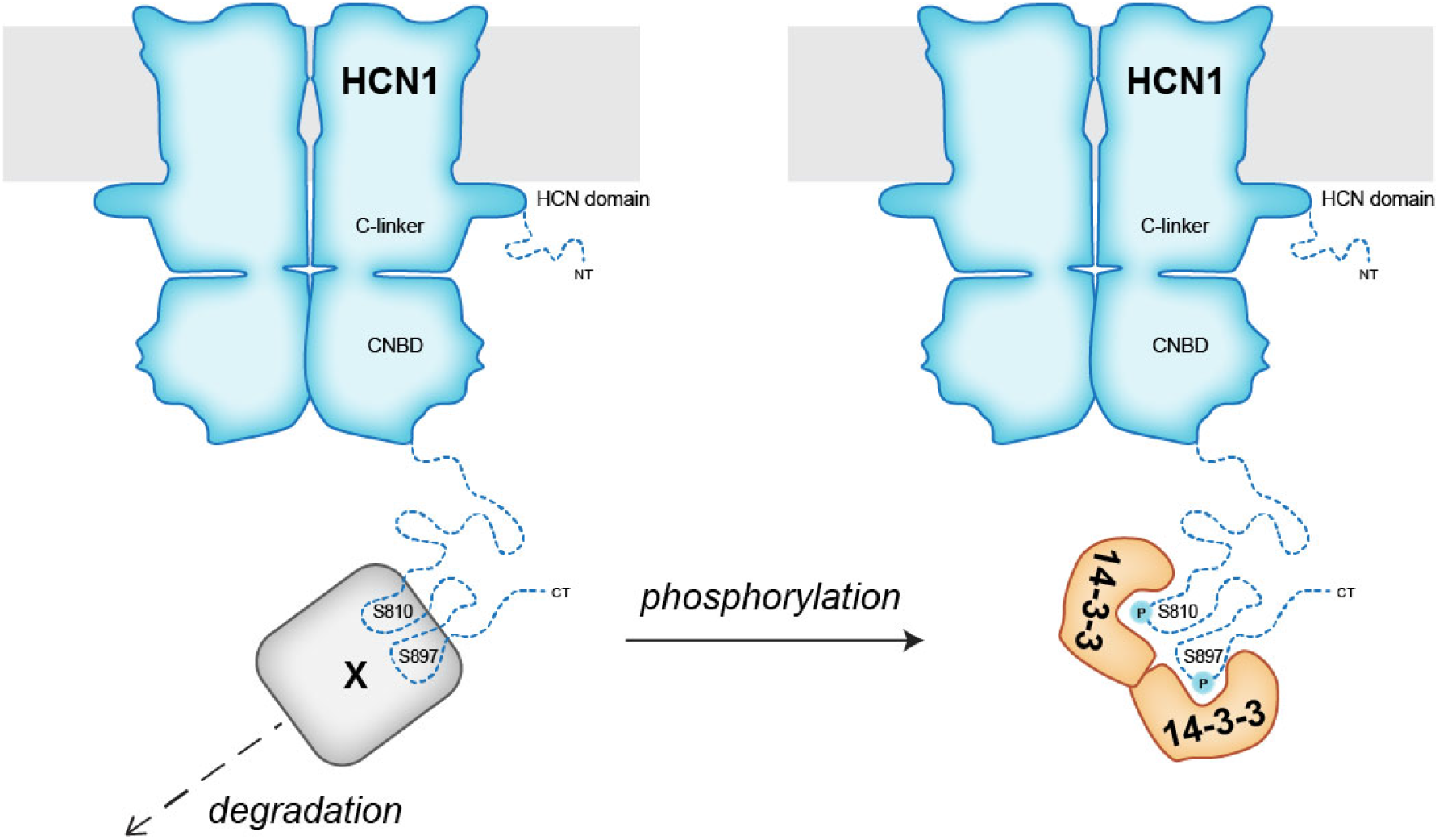
Model of proposed function of 14-3-3 binding to the distal C-terminus of HCN1. HCN1 degradation is induced by an unknown factor (X). HCN1 phosphorylation at S810 and S867 allows 14-3-3 binding, this blocks factor X from associating with HCN1 and prevents degradation of the channel.

## Discussion

The key finding of this study is the identification of HCN1 as a new client of the signaling adaptor 14-3-3. 14-3-3 can bind two sites, centered on S810 and S867 (corresponding to human residues S789 and S846), in the intrinsically disordered cytoplasmic C-terminus of mouse HCN1. While 14-3-3 can bind non-canonically to unphosphorylated sites and has recently been shown to bind O-GlcNAcylated-serine/threonine residues [56], our data support the conclusion that 14-3-3 binding to HCN1 is dependent on phosphorylation. This discovery is congruent with previous findings regarding HCN1 regulation. HCN1 phosphorylation has been implicated in either modulating channel activity or expression, likely depending on which sites are phosphorylated. Yet, the previously characterized HCN1 regulatory proteins are not known to require HCN1 phosphorylation for binding. In a recent phosphorylation site mapping study, one of the residues (S867) we identified as necessary for 14-3-3 binding was shown to be phosphorylated *in vivo*. In this study, peptides containing the second 14-3-3 binding site (S810) were not resolved, leaving it untested whether this site is phosphorylated *in vivo* [57]. While S867 phosphorylation was consistently identified in rat hippocampal CA1 tissue, there was no apparent change in the amount of S867 phosphorylation in a rat chronic epilepsy model [57]. Thus, when and why HCN1 S867 becomes phosphorylated in various neurons requires further investigation. We propose that along with S810, Ser867 phosphorylation represents a binding site for phosphorylation dependent modulation of HCN1 by 14-3-3.

We focused on the C-terminus of HCN1 as it contains the most evolutionarily conserved predicted 14-3-3 bindings sites but it is possible that 14-3-3 may bind to other regions of HCN1. The cytoplasmic N-terminus of HCN is intrinsically disordered, like the distal C-terminal tail, and contains several serine and threonine residues. This includes T39 which was shown to be phosphorylated [57] and lies within a high scoring predicted 14-3-3 binding site. 14-3-3 proteins function as dimers thus can bring different clients together or stabilize transient multi-protein complexes [58, 59]. For the sake of simplicity in our model we drew a single 14-3-3 dimer binding at the S810 and S867 sites in one HCN1 subunit. Alternatively, 14-3-3 could cross-link multiple subunits within one channel (or multiple channels) which would likely influence the stability of the channel. Different modes of 14-3-3 binding could elicit unique effects so it will be important to be mindful of these issues as future experiments move to testing the physiological effects of 14-3-3 mediated regulation in different neuronal populations.

One caveat of this study is the use of HEK293 cells as our primary model system. While a useful system to express the various constructs used in this study and examine the function of HCN1 mutants, we recognize that these cells do not adequately reflect neurons endogenously expressing HCN1. Thus, while the findings presented here implicate S810 and S867 as 14-3-3 binding sites, additional residues may be phosphorylated and accessible to 14-3-3 *in vivo*. The choice of model system likely explains the ambiguous results we obtained testing the effect of individual kinase inhibitors. That experiment would be better carried out *in vivo*, under physiologically relevant conditions, ideally where phosphorylation can be induced rather than relying on manipulation of any basal phosphorylation of HCN1. That said, it is intriguing that we obtained the strongest effects in HEK293 cells by inhibiting PKC and p38MAPK which have both been implicated in neuronal regulation of HCN1 [37, 60]. Another issue is that we do not know the ratio of basally phosphorylated to unphosphorylated HCN1 in HEK293 cells so it could well be that the unphosphorylated pool of HCN1 dominates which would make the serine to alanine substitutions effectively silent mutations. Phosphomimetic substitutions are often used to examine the function of site-specific phosphorylation; however, it is well characterized that negatively charged amino acids are generally insufficient to facilitate 14-3-3 binding, precluding our ability to use this strategy [48, 61-63].

The major conceptual challenge presented by our data was the observation that 10-amino acid deletions centered on S810 and S867 both prevented 14-3-3 binding and decreased the turnover rate of HCN1 leading to increased surface expression and higher current density. While site-specific serine-to-alanine mutations of S810 and S867 prevented 14-3-3 binding but did not significantly affect the turnover rate or expression of HCN1. One possibility is that 14-3-3 is a non-essential component of a complex that promotes HCN1 channel degradation. This hypothesis is unsatisfying and suggests that this interaction has no functional role despite S810 and S867 being highly conserved. A second possibility is that phosphorylation of S810 and S867 facilitates a domain recognition switch where the distal C-terminus is accessible to 14-3-3 when phosphorylated and 14-3-3 binding masks recognition by additional factors as modeled in Figure 8. This predicts that 14-3-3 binding would be protective and limit HCN1 degradation in a phosphorylation dependent manner. Consistent with this, we do see a slight reduction in the half-life of SNAP-HCN1_S810A; S867A_ (3.0hr vs 3.3hr for SNAP-HCN1).

The region containing the 14-3-3 binding sites is intrinsically disordered, so it is better to think of it as existing in a population of different conformations rather than in just one fixed state. That means binding by multiple proteins may not be competitive when HCN1 channels are considered en masse. Thus, we must consider how the various factors acting on the C-terminus of HCN1 act in coordination, whether they function in competitive or cooperative terms. This is especially important when we consider TRIP8b, an endogenous HCN channel accessory protein not present in HEK293 cells. TRIP8b binds to both the extreme C-terminal ser-asn-leu tripeptide of HCN1 and the cyclic nucleotide binding domain, spanning the intrinsically disordered region containing the 14-3-3 binding sites [29]. Depending on which splice isoform is expressed, this accessory subunit can either promote or inhibit expression of HCN1 such that TRIP8b may act in tandem or opposition with 14-3-3 [30-32]. Another HCN1 interacting partner that must be considered is Nedd4-2 which is a candidate for the unknown signaling factor promoting HCN1 degradation. Nedd4-2 is an E3 ubiquitin ligase known to target multiple ion channels including HCN1 [35, 64]. Nedd4-2 binds the C-terminus of HCN1, though it appears to bind to a more proximal region than 14-3-3. The lysine ubiquitinated by Nedd4-2 remains unknown and given the flexibility of the disordered C-terminus, may be at a more distal site in closer proximity to the 14-3-3 binding sites or may depend on the C-terminus adopting a conformation prevented by 14-3-3 binding. Testing of these models will be very challenging as it will be most informative if performed using *in vivo* systems that represent normal neuronal homeostasis and responsiveness to neuro-modulatory signaling.

Taken together, this work demonstrates that 14-3-3 binds the C-terminus of HCN1 in a phosphorylation dependent manner. Phosphorylation is implicated as a means to regulate HCN1 channel function and, to our knowledge, this is the first HCN1 interacting partner identified whose binding depends on phosphorylation of the channel. While we are unable to definitively identify the function of this interaction, our work demonstrates that 14-3-3 binding occurs at sites involved in regulating HCN1 turnover, identifying this region as an area of interest for further studies into regulation of HCN1 channels. Uncovering all the mechanisms regulating HCN1 is important as this channel is an attractive drug target for amelioration of various neurological and neuropsychiatric conditions.

## Experimental Procedures

### Molecular biology

eSpCas9(1.1)_No_FLAG_AAVS1_T2 was a gift from Yannick Doyon (Addgene plasmid # 79888; http://n2t.net/addgene:79888; RRID:Addgene_79888) [65]. The pSNAPf plasmid was obtained from New England Biolabs (cat. # N9183S) and the SNAP-tag was subcloned into a previously described plasmid encoding mouse HCN1 containing an HA-tag inserted into the extracellular loop between the S5 and S6 transmembrane domain (HA-HCN1) [55] to make SNAP-HCN1. pKanCMV-mRuby3-18aa-actin was a gift from Michael Lin (Addgene plasmid # 74255; http://n2t.net/addgene:74255; RRID:Addgene_74255) [66] and the mRuby3-tag was subcloned into the HA-HCN1 plasmid to make mRuby3-HCN1. While the HA-tag is not mentioned in this text, all HCN1 constructs used retain the HA-tag.

Homology directed repair templates for targeted knock in to the AAVS1 site were generated using AAVS1_Puro_PGK1_3xFLAG_Twin_Strep, a gift from Yannick Doyon (Addgene plasmid # 68375; http://n2t.net/addgene:68375; RRID:Addgene_68375) [65], as the base vector. SNAP-HCN1 was subcloned into this vector. In initial experiments SNAP-HCN1 expression was weak so we exchanged the PGK1 promotor for the CAG promotor from pCAG-mGFP, a gift from Connie Cepko (Addgene plasmid # 14757; http://n2t.net/addgene:14757; RRID:Addgene_14757) [67], to make AAVS1-CAG-SNAP-HCN1. The SNAP-tag was then exchanged for the mRuby3-tag from mRuby3-HCN1 to generate AAVS1-CAG-mRuby3-HCN1. 14-3-3 site mutations were made individually using site-directed mutagenesis and the double or triple mutant was then stitched together using in-fusion cloning to generate mRuby3-HCN1_Δ3, 14-3-3_ and SNAP-HCN1_Δ804-814; Δ861-871_. The cloning of serine to alanine substitutions into AAVS1-CAG-SNAP-HCN1 was performed by Genscript Cloning Services.

pDisplay-AP-CFP-TM was a gift from Alice Ting (Addgene plasmid # 20861; http://n2t.net/addgene:20861; RRID:Addgene_20861) [68]. The C-terminus of HCN1 was subcloned behind the transmembrane domain then the extracellular CFP tag was swapped for the SNAP-tag maying the SNAP-TM-HCN1_CT_ (CT_598-910_) reporter from which truncations were made up to the transmembrane domain (control reporter).

All sequences were validated by Sanger sequencing (Iowa Institute of Human Genetics, University of Iowa, Iowa City, IA or Eurofins Genomics, Louisville, KY). Novel plasmids are available upon request – AAVS1-CAG-mRuby3-HCN1 (wild-type and mutants), AAVS1-CAG-SANP-HCN1 (wild-type and mutants), and SNAP-TM-HCN1 (various truncations and targeted mutations).

### Cell culture

HEK293 (ATCC CRL-3216) were cultured in DMEM (Gibco cat. # 11965092) supplemented with 10% fetal bovine serum (Gibco cat. # 26140079), 100 U/mL penicillin, 100 µg/mL streptomycin (Gibco cat. # 15140122), and 0.5 µg/mL amphotericin B (Gibco cat. # 15290018). Cells were maintained at 37 °C with 5% CO_2_ in a humidifying incubator. Cells were passaged every two to three days and all experiments were performed prior to passage 25. Knock in cell lines were generated by co-transfecting cells 80% confluent in 6-well plates with 2 µg of AAVS1 homology directed repair template plasmid (wild-type and mutant versions of AAVS1-CAG-SNAP-HCN1 and AAVS1-CAG-mRuby3-HCN1) and 0.5 µg of AAVS1 targeted Cas9 plasmid (eSpCas9(1.1)_No_FLAG_AAVS1_T2). 24 hours after transfection, cells were treated with a high dose of puromycin, 5 µg/mL. After 48 hours, cells were switched to a maintenance dose of 2 µg/mL Puromycin which was maintained throughout. For transient transfections, 8 μg plasmid/10 cm plate or 2 μg plasmid/well of 6-well plate was used and cells harvested 24 hours post-transfection. All transfections were performed using Lipofectamine 2000 (ThermoFischer Scientific cat. # 11668027) at a 2:1 Lipofectamine (µL) to plasmid DNA (µg) ratio as described by the manufacturer.

### 14-3-3ζ purification

pPROEX-HTb vector for expression of human _6His_14-3-3ζ was a gift from Dr. Christian Ottmann [54]. This plasmid was transformed into BL21(DE3) *E. coli* and protein expression was induced with 1 mM isopropyl β-D-1-thiogalactopyranoside (IPTG) at mid-log phase. Induced cells were lysed using B-PER Bacterial Protein Extraction Reagent (ThermoFischer Scientific cat. # 90084) and purified using HisPur Ni-NTA resin (ThermoFisher Scientific cat. # 88221) or a HisTrap HP column (GE Healthcare). Purified _6His_14-3-3ζ was then dialyzed into PBS. Quality of the purification was assessed by SDS-PAGE and Coomassie staining, and protein concentration was measured by absorbance at 280nm.

### Isothermal titration calorimetry

ITC was performed as previously described [69]. Three phospho-peptides (A: ^653^SRMRTQ[pS]PPVY, B: ^804^VRPLSA[pS]QPSL, and C: ^861^TLFRQM[pS]SGAI) with numbering indicating position in mouse HCN1 were commercially synthesized (GenScript Peptide Synthesis Service). Experiments were conducted at 25 °C in PBS (pH 7.4) using an iTC200 (MicroCal). The concentration of _6His_14-3-3ζ in the cell was 70, 60, or 43 μM, while the concentration of peptides in the syringe was 700, 600, or 430 μM for phospho-peptides A, B, and C respectively; 20 injections of peptide were made per experiment. All experiments were performed in at least duplicate. The titration curve was fit to determine the binding stoichiometry, dissociation constant (K_D_), and thermodynamic parameters (enthalpy and entropy changes) using the Origin software provided by the manufacturer. ITC data is presented at mean ± standard deviation across replicates.

### Electrophysiology

Current recordings from HEK293 cell lines expressing mRuby3-HCN1 or mRuby3-HCN1_Δ3, 14-3-3_ were performed by Eurofins Discovery (St Charles, MO, USA). Recordings were made using the Qpatch automated patch clamp system. Current recordings at tests pulses from –30 to –130 mV were provided, as was the capacitance of each cell. From the data provided, the current density and percentage of maximal current were obtained and plotted against the voltage. Plots were fit to a Boltzman sigmoidal. All electrophysiology data is presented as mean ± standard error. For comparisons between cell lines, the maximal current density was compared using t-test.

### Antibodies and western blotting

All antibodies used are detailed in Table 2 and western blotting was performed using a standard protocol. Premade 4-20% TGX gels with either the 10-well 50 μL or 15-well 15 μL formats (Bio-Rad cat. # 4561094 and 4561096) were used in all instances. Sample buffer was prepared using 4X LDS buffer (ThermoFischer Scientific cat. # NP0007) and 10X reducing agent (ThermoFischer Scientific cat. # NP0009). SDS-PAGE was performed with a tris-glycine running buffer (25 mM Tris, pH 8.3, 192 mM glycine, 0.1% SDS). Transfer was then performed with a tris-glycine transfer buffer (25 mM Tris, pH 8.3, 192 mM glycine, 15% methanol, 0.01% SDS). Proteins were transferred to low fluorescence PVDF membrane (Millpore Sigma cat. # IPFL00010). REVERT total protein stain was used to detect and quantify transferred proteins. Prior to immunolabeling, membranes were blocked with Intercept blocking buffer (LI-COR cat. # 927-60001). Primary antibody was diluted in a 50/50 mixture of blocker and wash buffer (39 mM Tris, pH 7.5, 300mM NaCl, 0.1% Tween-20) and incubated on membrane overnight at 4°C. Membranes were washed with wash buffer prior to and after 1 hour secondary antibody incubation also diluted in a 50/50 mixture of blocker and wash buffer. For co-immunoprecipitation experiments blots were imaged using the ImageQuant LAS 4000 (GE Healthcare). Image adjustments to brightness and contrast were made using Adobe PhotoShop 2020. All other blots were imaged using a LI-COR Odyssey Fc imager and images were quantified and edited using the Image studio software (Ver 5).

**Table 2:** Antibodies

| Table 2: Antibodies |  |  |  |
| --- | --- | --- | --- |
| Antibody | Reference | Application | Figure |
| rabbit anti-HCN1 | Pan et al., 2014 [70] | western blotting against HCN1 in co-IP and pulldown experiments | 4 |
| mouse anti-HCN1 | UC Davis/NIH NeuroMab Facility Cat# N70/28, RRID:AB_2877279 | western blotting against mRuby3-HCN1 | 3 |
| rabbit anti-14-3-3 | Santa Cruz Biotechnology Cat# sc-629, RRID:AB_2273154 | western blotting against 14-3-3 in co-IP experiments | 3A |
| mouse anti-14-3-3 | Thermo Fisher Scientific Cat# MA5-12242, RRID:AB_10986852 | Immunoprecipitating antibody for co-immunoprecipitation experiments | 3A |
| mouse anti-GFP | Takara Bio Cat# 632380, RRID:AB_10013427 | Control antibody for co-immunoprecipitation experiments | 3A |
| rabbit anti-SNAP | New England Biolabs Cat# P9310S, RRID:AB_10631145 | western blotting against SNAP-TM reporters and SNAP-HCN1 | 5, 6, and 7 |
| mouse anti-NKA | Santa Cruz Biotechnology Cat# sc-58628, RRID:AB_781525 | western blotting in surface biotinylation experiments | 3D and 7A |
| mouse anti-GAPDH | DSHB Cat# DSHB-hGAPDH-2G7, RRID:AB_2617426 | western blotting in surface biotinylation experiments | 3D and 7A |
| TruBlot anti-mouse-HRP | Rockland Cat# 18-8817-30, RRID:AB_2610849 | secondary antibody used in co-IP experiments | 4A |
| goat anti-mouse800 | LI-COR Biosciences Cat# 925-68071, RRID:AB_2721181 | secondary antibody |  |
| goat anti-rabbit680 | LI-COR Biosciences Cat# 925-32210, RRID:AB_2687825 | secondary antibody |  |
| goat anti-mouse800 | Thermo Fisher Scientific Cat# SA5-35521, RRID:AB_2556774 | secondary antibody |  |
| goat ant-rabbit 680 | Thermo Fisher Scientific Cat# 35568, RRID:AB_614946 | secondary antibody |  |

### Co-immunoprecipitation and pulldown assays

All samples were homogenized on ice in lysis buffer (1% Triton X-100 in PBS: 137mM NaCl, 2.7 mM KCl, 10 mM Na_2_PO_4_, 1.8 mM KH_2_PO_4_, pH 7.4) supplemented with cOmplete Mini, EDTA free protease inhibitor cocktail and PhosSTOP phosphatase inhibitor cocktail (Millipore Sigma cat. # 11836170001 and 4906845001). For tissue preparation, wildtype (C57Bl6/J) or HCN1 knockout mice as described in [70] were used. Whole brain was removed and homogenized in 5 mL of lysis buffer and 4-6 retina were homogenized in 1 mL of lysis buffer. HEK 293 cells were harvested by addition of lysis buffer directly to the culture plate. For full-length HCN1, transfected HEK293 cells on a 10 cm plate were lysed with 1 mL of lysis buffer. For the SNAP-TM reporters, transfected HEK293 cells on 6-well plates were lysed with 400 µL of lysis buffer per well. BCA assays (ThermoFischer Scientific cat. # 23227) were used to measure protein concentration.

For the 14-3-3 immunoprecipitation, 2.5 μg of anti-14-3-3 or anti-GFP antibodies were immobilized on 20 μL of protein A/G for two hours at 4 °C. Equal amounts of lysate were added to antibody-bound resin overnight at 4°C and washed with lysis buffer. Protein was eluted with 2X reducing sample buffer.

For 14-3-3 pulldowns, _6His_14-3-3ζ was incubated with Ni-NTA resin in 5% BSA dissolved in lysis buffer supplemented with 25 mM imidazole for 3 hours at 4°C; 5 mg of _6His_14-3-3ζ was used for pulldown from brain, 0.5 mg for pulldown from retina and HEK293 cells expressing full-length HCN1, 0.1 mg for pulldown of SNAP-TM-HCN1_CT_ reporters. For the control, Ni-NTA resin was treated identically without the addition of _6His_14-3-3ζ. Equal amounts of lysate were added to control or _6His_14-3-3ζ conjugated resin and incubated overnight at 4 °C. Resin was then washed with lysis buffer supplemented with 25 mM imidazole and protein was eluted with 250 mM imidazole in 8% SDS in 0.1 M Tris pH 7.8.

For pulldowns using the SNAP-TM reporters, the amount of each reporter pulled down was compared using a t-test or 1-way ANOVA followed by Dunnett’s multiple comparison to compare the truncated or mutant reporters to control (CT_598-910_ or CT_731-910_). All pulldown data presented as mean ± standard deviation.

### Surface Biotinylation

HEK293 cells were briefly washed with cold PBS prior to incubation with 2 mg/mL Sulfo-NHS-SS-Biotin (APExBIO cat. # A8005) in cold PBS for thirty minutes on ice. The reaction was quenched by the addition 50 mM Tris-HCl, 150 mM NaCl, pH 7.5). Cells were briefly washed with PBS prior to addition of lysis buffer. For each replicate, equal amounts of protein (200-300 µg) for each experimental sample was added to 75 μL of High capacity Neutravidin agarose (ThermoFischer Scientific cat. # 29202) and incubated at 4°C overnight. Resin was washed with lysis buffer supplemented with 0.5% SDS. Protein was eluted by incubation for 5 minutes at 75°C in 2X reducing sample buffer prepared from 4X LDS sample buffer and 10X reducing agent. Eluted protein was then examined by western blot. T-test was used to compare mRuby3-HCN1 and mRuby3-HCN1_Δ3, 14-3-3_. 1-way AVONA followed by Dunnett’s multiple comparison was used to compare SNAP-HCN1_Δ804-814; Δ861-871_ and SNAP-HCN1_S810A; S867A_ to SNAP-HCN1. All data presented as mean ± standard deviation.

### Pulse-chase experiments

Stable HEK293 cell lines were grown to confluency on a 35mm dish. The day before the pulse-chase experiment, cells were trypsinzied, resuspended in 10 mL of media, and 0.5 mL of the cell suspension was plated in each well of a 24-well plate. The following day cells were treated with 1 µM SNAP-cell TMR-Star (New England BioLabs cat. # S9105S) in complete culture media for 45 minutes at 37 °C; for the chase, media was replaced with media containing 3 µM SNAP-Cell block (New England BioLabs cat. # S9106S) and cells maintained at 37 °C. Aliquots of cells were harvested immediately at the end of the pulse or at 4, 8, 12, and 24 hours into the chase. Lysates were immediately frozen and processed once the final timepoint was collected. Lysates were quantified by BCA assay and equal total protein concentrations were analyzed by western blotting against the SNAP-tag with membranes imaged for SNAP-HCN1 conjugated SNAP-Cell TMR-star (600 nm channel on LI-COR Odyssey FC) prior to blocking.

The labeled SNAP-HCN1 fluorescent signal was normalized to SNAP-HCN1 western blot intensity. This value was then normalized to the 0-hour chase timepoint to give the % labeled SNAP-HCN1. The half-life was calculated by fitting the data to a one-phase decay curve. 2-way ANOVA followed by Dunnett’s multiple comparison was used to compare SNAP-HCN1_Δ804-814; Δ861-871_ and SNAP-HCN1_S810A; S867A_ to SNAP-HCN1. Data presented as mean ± standard deviation.

## Data availability

Detailed descriptive statistics for each experiment are available on request from the corresponding author.

## Supporting information

*This article contains supporting information; a table of kinase inhibitors used to generate Figure 6*.

## Funding and additional information

This work was supported by the National Eye Institute (R01 EY020542 to SAB). The content is solely the responsibility of the authors and does not necessarily represent the official views of the National Institutes of Health.

## Abbreviations and nomenclature

(HCN1): The Hyperpolarization activated cyclic nucleotide gated channel 1
(ITC): isothermal titration calorimetry
(NKA): Na^+^/K^+^-ATPase
(GAPDH): glyceraldehyde 3-phosphate dehydrogenase
(CNBD): Cyclic Nucleotide Binding Domain.

## Figures and Figure legends

**Table S1:**
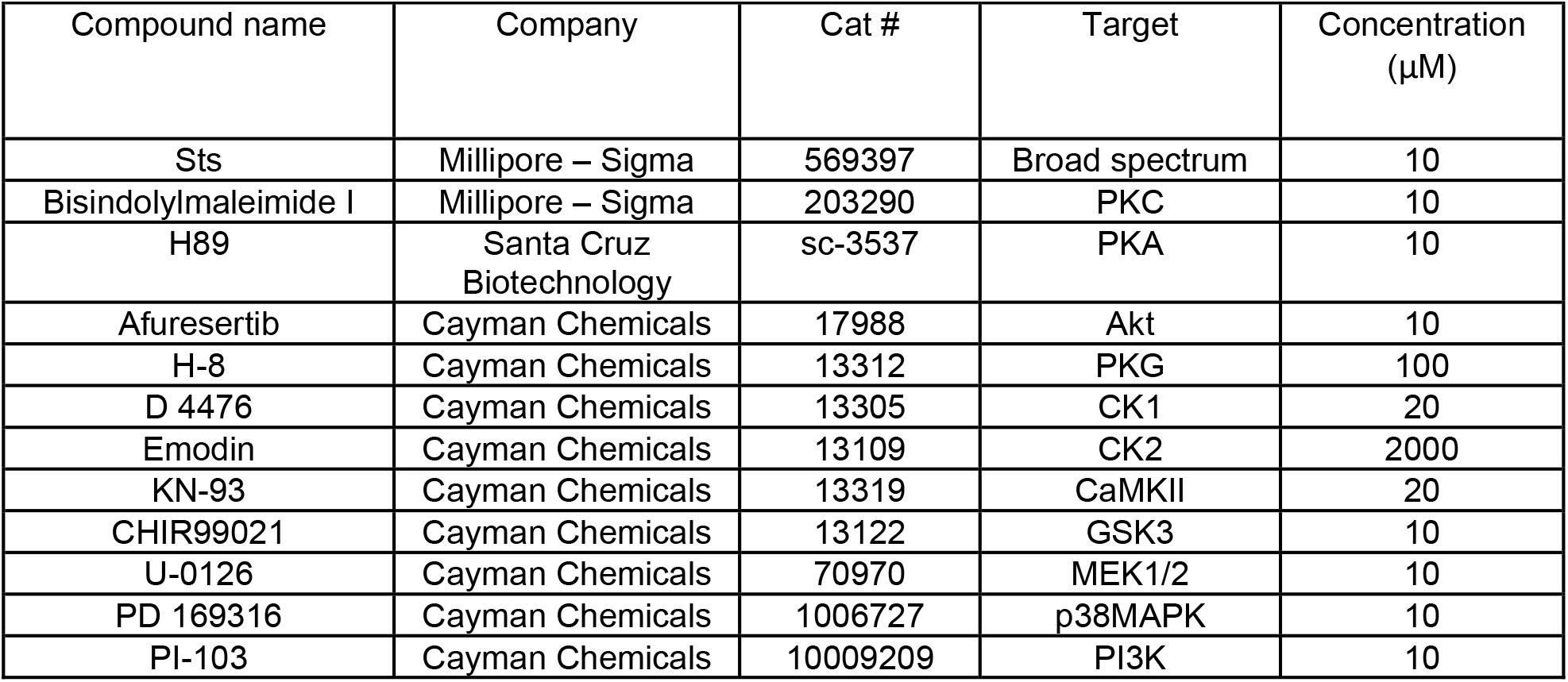
Kinase inhibitors used in Figure 6

| Compound name | Company | Cat # | Target | Concentration (μM) |
| --- | --- | --- | --- | --- |
| Sts | Millipore – Sigma | 569397 | Broad spectrum | 10 |
| Bisindolylmaleimide I | Millipore – Sigma | 203290 | PKC | 10 |
| H89 | Santa Cruz Biotechnology | sc-3537 | PKA | 10 |
| Afuresertib | Cayman Chemicals | 17988 | Akt | 10 |
| H-8 | Cayman Chemicals | 13312 | PKG | 100 |
| D 4476 | Cayman Chemicals | 13305 | CK1 | 20 |
| Emodin | Cayman Chemicals | 13109 | CK2 | 2000 |
| KN-93 | Cayman Chemicals | 13319 | CaMKII | 20 |
| CHIR99021 | Cayman Chemicals | 13122 | GSK3 | 10 |
| U-0126 | Cayman Chemicals | 70970 | MEK1/2 | 10 |
| PD 169316 | Cayman Chemicals | 1006727 | p38MAPK | 10 |
| PI-103 | Cayman Chemicals | 10009209 | PI3K | 10 |

